# Evolutionary adaptation of helminth immunomodulators to their hosts

**DOI:** 10.64898/2026.09.02.748793

**Authors:** Beatrice Ward, Abhishek Jamwal, Florent Colomb, Adefunke Ogunkanbi, Olivia C A Fleming, Sophie Darroch, Findlay A J Donnelly, Lucy Gregory, Danielle J Smyth, Amy B Pedersen, Lewis Stevens, E Suzanne Cohen, Matthew K Higgins, Henry J McSorley

**Author notes:** Corresponding author: Henry J McSorley. Email address of all authors.

## Abstract

**Background:** Parasitic nematodes have evolved secreted effector proteins which interact with host molecules to modulate the host immune response. These interactions can be highly specific and are under strong selective pressure. Here, we investigate the host-pathogen interaction between *Heligmosomoides bakeri* and its natural host *Mus musculus*, and between *H. polygyrus* and its natural host *Apodemus sylvaticus*. We focus on the Alarmin Release Inhibitor family (previously called HpARI) produced by heligmosomid parasites, which interact with IL-33 and heparan sulphate.

**Results:** Using comparative genomics, we identified duplications and diversifications of the ARI family including multiple copies of ARI1 and conserved ARI3 homologues in both *H. bakeri and H. polygyrus.* We identify a new family member, HpolARI4, encoded by *H. polygyrus,* as well as a more distant homologue, HmixARI, encoded by *Heligmosomum mixtum,* a parasite of the bank vole (*Myodes glareolus).* Our functional assays identify that HpolARI4 preferentially blocks *A. sylvaticus* IL-33 but has little activity against *M. musculus* IL-33, whereas HbakARI2 is highly specific for *M. musculus* IL-33. We furthermore show differential interaction with host heparan sulphate, with HbakARI2 and HpolARI4 showing high affinity binding, whereas HbakARI3 shows no binding. We further test this family against non-host human IL-33 and show that functional suppression can be achieved by HbakARI3 *in vitro* and *in vivo*.

**Conclusion:** By integrating genomic data, structural insights, and functional assays, we reveal how the expansion and diversification of the ARI family drive host-specific IL-33 modulation, providing a molecular model for nematode-host co-evolution.

## Introduction

The host-parasite relationship is defined by co-evolution, in which parasites evolve to better infect, survive, and replicate within their hosts, while hosts evolve to develop better defences to reduce damage and limit infection, thereby driving further parasite evolution. This co-evolutionary dynamic is often referred to as the Red Queen hypothesis (1) and has led to a framework proposing that genes involved in these interactions undergo rapid evolution in both host and parasite (2). This interaction includes some evolutionary trade-offs, as parasite proteins involved in interactions with the host (such as receptor-binding domains, or immune evasion proteins) are under strong diversifying selection for mutations that reduce antibody recognition (3), while also being under positive selection to maintain strong binding to host protein targets. Conversely, the host’s target proteins are under diversifying selection to escape parasite recognition or immunomodulation, while maintaining their function within the host. While this interaction is well understood in viruses, bacteria, and single-celled parasites (where pathogens often use host proteins as cellular entry receptors), little is understood about how, at a molecular level, nematode parasites co-evolve with their hosts (2).

Parasitic nematodes are a major global health burden causing widespread morbidity in humans, and result in expensive treatments to control spread within livestock (4, 5). While host specificity of nematode parasites is well-described (6, 7), the molecular basis of this specificity is not well understood. Due to their size, nematode parasites cannot invade host cells and instead infect their hosts through largely mechanical processes, involving physical invasion as well as secretion of proteases to degrade host barriers (8). Therefore, unlike viruses, bacteria, and unicellular parasites, nematode parasites do not have defined protein receptor-ligand interactions that govern infectivity and can be studied to characterise these co-evolutionary processes. However, parasitic nematodes secrete immunomodulatory proteins, some of which bind to host immune proteins, providing a basis for molecular evaluation of the host-pathogen molecular interface (8).

*Heligmosomoides* parasites are intestinal nematodes that infect various species of rodents. *Heligmosomoides* nomenclature has been inconsistent; in studies comparing infections in wood mice (*Apodemus sylvaticus*) versus laboratory isolates, the parasite that infects *A. sylvaticus* is named *Heligmosomoides polygyrus,* while the parasite that infects *Mus musculus* is named *Heligmosomoides bakeri* (9). Meanwhile in the immunological literature, *Heligmosomoides* parasites used to infect *M. musculus* in the laboratory are often referred to as *H. polygyrus* or *H. polygyrus bakeri* (10). Recent genomic comparisons reveal a surprising level of divergence between the two species, as well as extensive diversity within populations of each species (11). Although both *Heligmosomoides* species can infect both hosts, much reduced survival and chronicity of infection in mismatched host-parasite infections suggests evolutionary adaptation of each parasite to their preferential host (12). The heligmosomid family also contains more distant members including *Heligmosomum mixtum* and *Heligmosomoides glareoli,* which are natural parasites of the bank vole *Myodes glareolus* and which, like *H. polygyrus* and *H. bakeri*, are proposed to secrete immunomodulatory factors (13).

Of the secreted immunomodulators of *Heligmosomoides* parasites, we previously discovered the Alarmin Release Inhibitor family of proteins. In our previous work, we referred to these proteins, identified in *H. polygyrus bakeri* as “HpARI1”, “HpARI2” and “HpARI3” (14). Here, to differentiate proteins from different heligmosomid parasites, we will refer to the parasite as “*H. bakeri*” and the immunomodulators as “HbakARI” to allow comparison between different parasite species. Members of this family of proteins each consist of 3 complement control protein (CCP) domains and can bind two partners: the CCP1 domain binds to heparan sulphate (HS; a highly abundant glycosaminoglycan of the extracellular matrix (15)), and the CCP2 and CCP3 domains together bind to host IL-33. The efficacy of these functions differs between family members: while HbakARI1 and HbakARI2 bind strongly to *M. musculus* cytokine IL-33 and effectively suppress responses to it (16), HbakARI3 binds but does not block responses to the cytokine (14). Furthermore, while HbakARI2 binds strongly to HS, HbakARI1 has reduced binding efficacy, while HbakARI3 shows no binding (17). Lastly, in *M. musculus* infection with *H. bakeri*, only vaccination with HbakARI2 provided protection against infection, while HbakARI1 or HbakARI3 vaccination had no effect (18). Therefore, despite good conservation of sequence between these ARI proteins, their effects differ greatly *in vitro* and *in vivo*.

The availability of a structure of HbakARI2 bound to *M. musculus* IL-33 (16) now enables us to better understand the host-pathogen coevolutionary interface in atomic detail. Furthermore, the role of IL-33 in type 2 immunity and human immune-mediated diseases (19) has led to our proposal (20) for the potential use of ARI proteins as therapeutic agents in humans. However, humans do not harbour *Heligmosomoides* parasites; therefore, humans are not a species that *Heligmosomoides* has co-evolved with. To investigate the adaptation of the ARI family to its hosts, and to translate these findings to humans, we therefore tested each of the known HbakARIs against *M. musculus*, *A. sylvaticus,* and *Homo sapiens* IL-33. Furthermore, we identify a new member of the ARI family (HpolARI4) in the *H. polygyrus* genome, and a more distant homologue in the *H. mixtum* genome (HmixARI). We find that the ARI family shows stark differences in their affinity and suppressive effects against IL-33 from different host species, and efficacy of binding to host HS. Thus, despite relatively small sequence divergences, this family of immunomodulators shows specificity towards different targets, and represents an evolutionary adaptation to their hosts.

## Results

### HpARI2 and HpARI3 suppress human IL-33 with different efficacies

We first investigated the effects of *H. bakeri* ARI proteins on non-host (human) IL-33, using transgenic mice in which the *M. musculus* IL-33 gene had been replaced with the human IL-33 gene (“hIL-33tg” mice), resulting in human IL-33 release on allergen stimulation (21). We administered Alternaria allergen as an IL-33 stimulus, co-administered with HbakARI1, HbakARI2 or HbakARI3 recombinant proteins, and collected bronchoalveolar lavage (BAL) 30 min later to assess levels of human IL-33 released. While HbakARI1 had no effect, HbakARI2 and HbakARI3 both suppressed IL-33 detection in these BAL samples (**Figure 1A**). This indicates that HbakARI2 and HbakARI3 (but not HbakARI1) bind hIL-33 *in vivo*.

**Figure 1:**
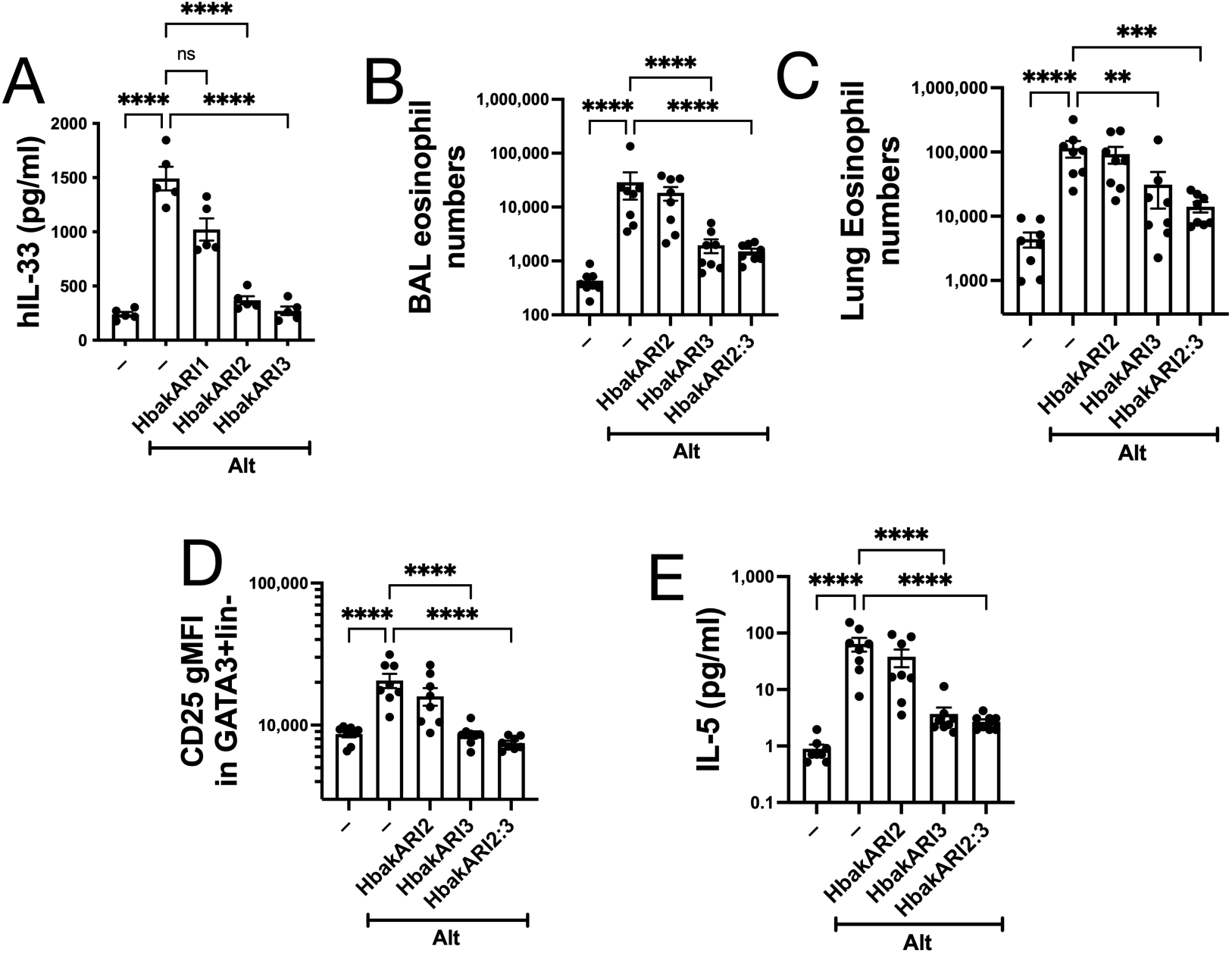
HbakARI3 shows suppressive capacity against human IL-33. Alternaria and HbakARH, HbakARI2 or HbakARI3 were intranasally administered to hlL-33tg mice, which were culled 30 min later to assess human IL-33 levels in bronchoalveolar lavage by ELISA **(A).** Alternaria and HbakARI2, HbakARI3 or HbakARI2:3 fusion were intranasally administered to humanized IL-33 mice, which were culled 24 h later. Flow cytometry was used to assess eosinophils numbers in bronchoalveolar lavage **(B)** and lung tissue **(C),** and CD25 geometric mean fluorescence intensity on lung type 2 innate lymphoid cells **(D).** IL-5 in bronchoalveolar lavage supernatants was measured by ELISA **(E).** **(A)** shows n=5 per group, representative of 2 repeat experiments, **(B-E)** are pooled from 2 repeat experiments, for a total n=8 per group. Standard error of mean shown. Data analysed by one way ANOVA comparing all groups to positive control, ** = p<0.01, *** = p<0.001, **** = p<0.0001.

Our previous studies with HbakARI3 indicated that binding to IL-33 can result in steric hindrance of antibodies used to detect the cytokine by ELISA but does not always correlate with blockade of the downstream effects of IL-33 (14). Therefore, the hIL-33tg mouse experiment was repeated, and downstream responses to released hIL-33 (which can signal through the mouse IL-33 receptor (21)) were measured 24 h later. Furthermore, we also applied an HbakARI2:3 fusion protein, which consists of the HS-binding domain of HbakARI2 (domain 1) fused to the IL-33-binding domains of HbakARI3 (domains 2 and 3) (17). When assessing BAL and lung eosinophilia, lung ILC2 CD25 expression (a measure of ILC2 activation) and BAL IL-5 levels, HbakARI2 was ineffective in suppressing the response to hIL-33 release *in vivo*. Both the HbakARI3 and the HbakARI2:3 fusion proteins were highly effective in blocking these responses (**Figure 1B-E**), indicating that the IL-33-binding domains were solely responsible for this activity.

These data, combined with our previous studies (14), indicate that members of the ARI family can have differential effects on IL-33 from different host species. We therefore decided to investigate the conservation of the ARI family across *Heligmosomoides* species, and the effects of the ARIs on other host species.

### Identification of HpolARI4, a new member of the HpARI family

HbakARI1, HbakARI2 and HbakARI3 were identified from previous genomic and transcriptomic studies of *H. bakeri* (22). Recently, improved chromosome-level genomes of *H. bakeri* (from *M. musculus* infections) and *H. polygyrus* (from *A. sylvaticus* infections) were published (11), allowing us to identify ARI homologues in both genomes (**Table 1, Supplementary Figure 1**).

**Table 1:** ARI family members in *H. bakeri* and *H. polygyrus* genomes.

| ARI | Species | Position in genome and strand | Ref annotation |
| --- | --- | --- | --- |
| ARI1 | *H. bakeri* | 1. Chromosome V 3,178,410:3,181,022 minus<br>2. Chromosome V 3,192,364:3,194,976 minus<br>3. Chromosome V 3,206,324:3,208,934 minus<br>4. Chromosome V 3,220,309:3,222,919 minus<br>5. Chromosome V 3,234,272:3,236,882 minus<br>6. Chromosome V 3,248,225:3,250,558 minus | 1. Not annotated<br>2. Not annotated<br>3. Not annotated<br>4. Not annotated<br>5. Not annotated<br>6. Not annotated |
| ARI1 | *H. polygyrus* | 1. Chromosome V 3,214,755:3,217,373 plus<br>2. Chromosome V 3,237,870:3,240,488 minus<br>3. Chromosome V 3,260,544:3,263,163 minus<br>4. Chromosome V 8,620,460:8,623,075 plus | 1. ngHelPoly1.g13325<br>2. ngHelPoly1.g13326<br>3. ngHelPoly1.g13327<br>4. ngHelPoly1.g13481 |
| ARI2 | *H. bakeri* | 1. Chromosome III 106,672,246:106,674,807 minus | 1. nxHelBake1.g18769 |
| ARI2 | *H. polygyrus* | Not found | – |
| ARI3 | *H. bakeri* | 1. Chromosome III 106,829,021:106,831,558 minus<br>2. Chromosome I:96,076,654-96,131,422 minus | 1. Not annotated<br>2. nxHelBake1.g12877 |
| ARI3 | *H. polygyrus* | 1. *Isotig* CANDYH010000565 9,031:11,622 plus | 1. ngHelPoly1.g20441 |
| ARI4 | *H. bakeri* | Not found | – |
| ARI4 | *H. polygyrus* | 1. Chromosome III 103,758,595:103,761,159 minus | 1. ngHelPoly1.g13087 |

**Supplementary Figure 1:**
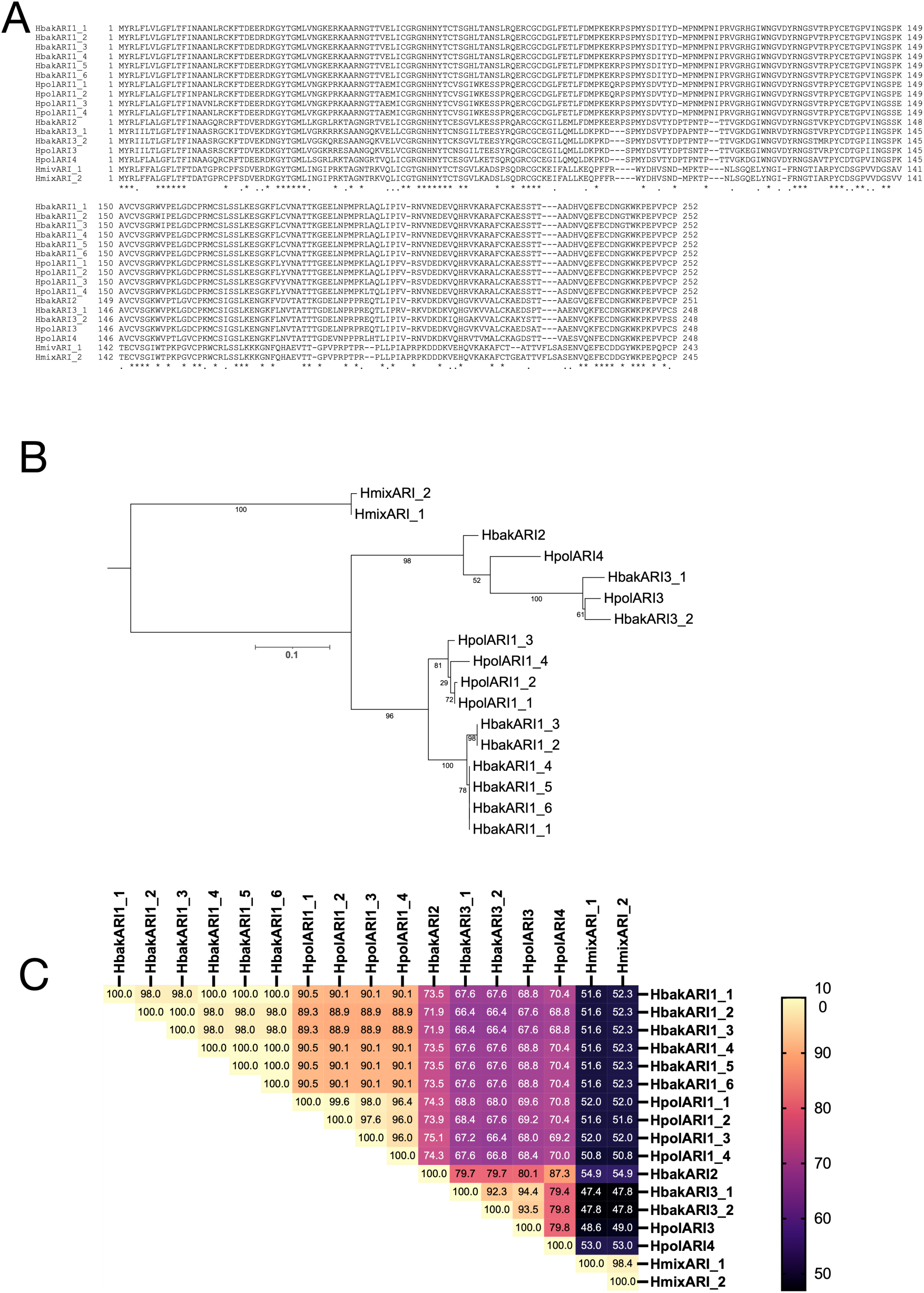
ARI homologues in *H. bakeri, H. polygyrus* and *H. mixtum.* Sequence alignment of ARI family members from *H. bakeri* and *H. polygyrus* (identified in Table 1), with ARI homologues from *H. mixtum* (A). Phylogenetic tree of all ARI family members (B). Matrix showing % identity of each ARI homologue.

In the *H. bakeri* genome, six copies of genes encoding HbakARI1 were found clustered on chromosome V. The six *H. bakeri* HbakARI1 genes show very high sequence similarity to each other, with the first 5 copies showing >99% nucleotide identity across the entire gene, and the sixth copy showing slightly lower (84%) identity. Predicted HbakARI1 proteins were 98-100% identical (i.e. 0-5 amino acid polymorphisms across the full sequence of these proteins) (**Supplementary Figure 1C**). Furthermore, each copy is encoded in the same orientation (minus strand) and is regularly spaced in the genome, with an 11.3 kb interval between every copy (**Figure 2A**). The high degree of sequence identity of these genes and surrounding sequence, as indicated by multiple dark blue parallel diagonal lines in Figure 2A, suggests they have undergone recent tandem duplication.

**Figure 2:**
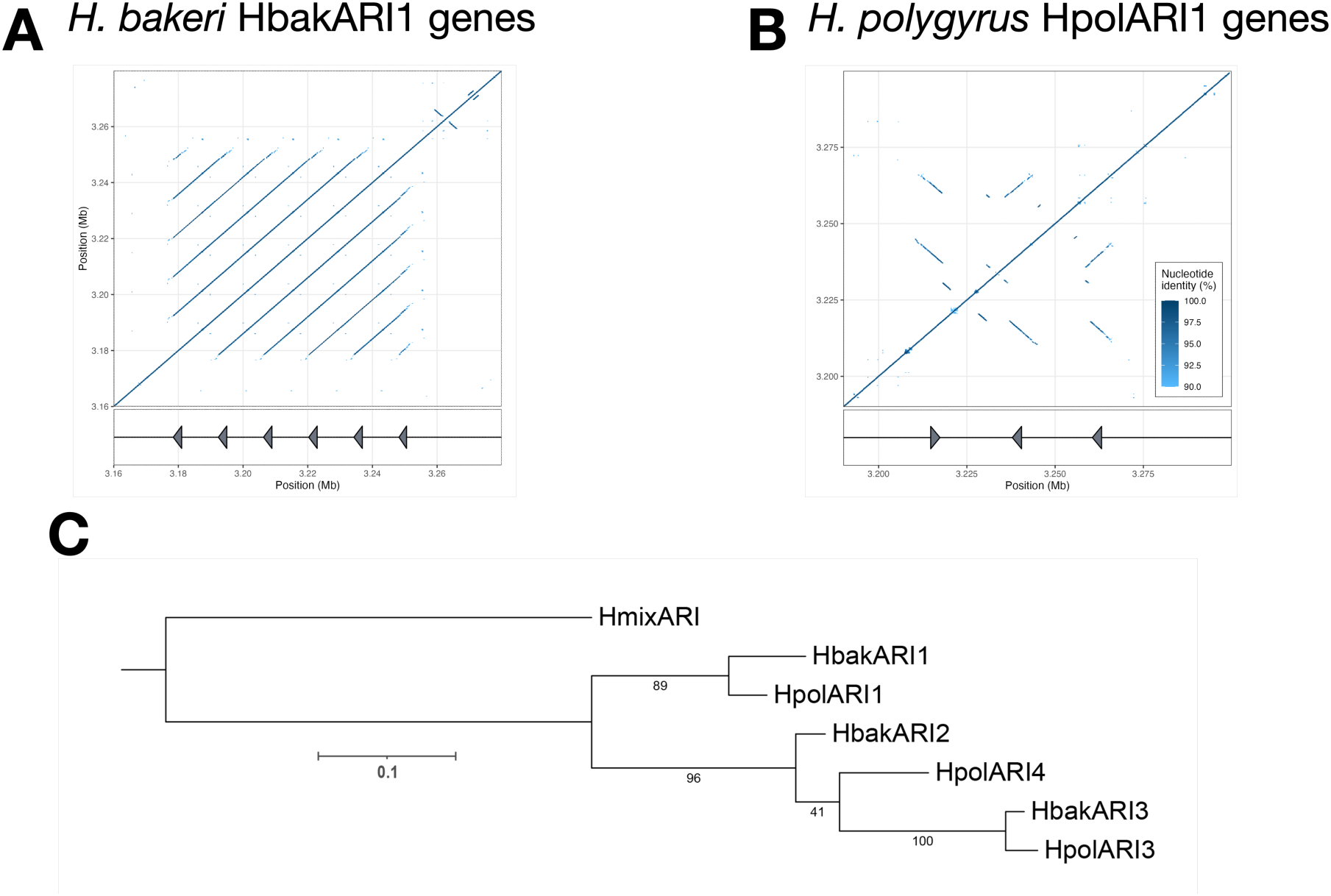
Dot plot ARI1 gene copies, showing similarity and spacing in *H. bakari* (A) and *H. polygyrus* (B) genomes. HpolARI1-4 is not shown due to distance from other HpolARI1 copies. Phylogenetic tree of ARI consensus sequences, roted using the *H. mixtum* HmixARi1 sequence as an outgroup (C). Bootstrap support values are shown. Branch lengths represent the number of substitutions per site; scale is shown.

**Supplementary Figure 2:**
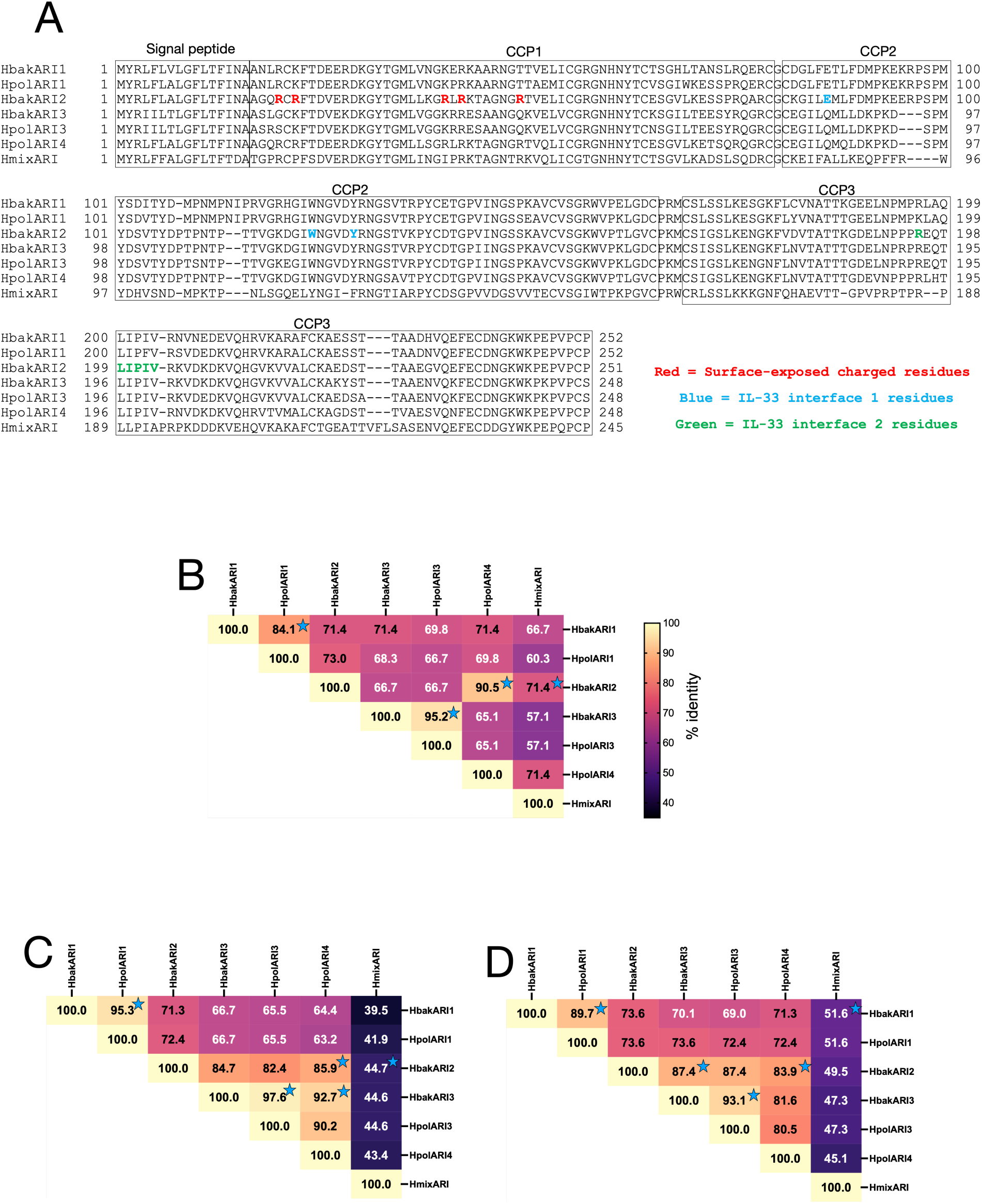
ARI homologue CCP domain identities s in *H. bakeri, H. polygyrus* and *H. mixtum.* (A) Sequence alignment of ARI homologue sequences from *H. bakeri, H. polygyrus* and *H. mixtum.* Critical residues involved in surface and IL-33 binding identified from HbakARI2 structure are indicated. Percentage identity matrices comparing the (B) CCP1, (C) CCP2, and (D) CCP3 domains. Blue stars indicate highest identity pairs for each CCP domain, showing that *H. bakeri* and *H. polygyrus* ARI1s and ARI3s CCP domains are most similar to each other in each case, while HpolARI4 CCP1 is most similar to HbakARI2 CCP1, HpolARI4 CCP2 is most similar to HbakARI3, and HpolARI4 CCP3 is most similar to HbakARI2.

On chromosome V of the *H. polygyrus* genome, there were 4 copies of a gene with ∼90% predicted amino acid identity to HbakARI1, which we identified as HpolARI1. The HpolARI1 genes showed slightly lower similarity over the whole gene compared to the HbakARI1 copies, ranging from 95.7-99.7% nucleotide identity and with >96% identity at the protein level. However, unlike *H. bakeri*, the *H. polygyrus* HpolARI1 genes were not all regularly spaced in the genome (copies 1-3 separated by ∼20 kb, copy 4 around 5.4 Mb distant), and while copies 2 and 3 were on the minus strand, copies 1 and 4 were on the plus strand (**Figure 2B**). The level of similarity (including in non-coding regions), orientation, and spacing indicates that the ARI1 family was duplicated and diversified more recently in *H. bakeri* than in the *H. polygyrus* genome. Moreover, phylogenetic analysis of the HbakARI1 and HpolARI1 genes suggests that the duplications occurred independently after the two species diverged (**Figure 2C**).

In the *H. bakeri* genome, a single copy of the HbakARI2 gene was found on chromosome III, while 2 copies of genes encoding close HbakARI3 homologues were found on chromosome III and I respectively (**Table 1**). In *H. polygyrus*, a gene encoding a protein with >90% identity to HbakARI3 was identified in a non-chromosomal contig. However, no genes were identified in the *H. polygyrus* genome with >90% identity to HbakARI2. Instead, a gene encoded at a similar locus on chromosome III was identified with 87% identity for HbakARI2, but which also contained conserved sequence elements in the CCP2 domain with HbakARI3 and HpolARI3 (**Supplementary Figure 2**). This new member of the HpARI family was named HpolARI4.

Finally, two more distant ARI homologues were identified by homology searching of the recently-published *H. mixtum* genome (23). These *H. mixtum* ARI homologues were 98% identical to each other, but formed a clear outgroup compared to the other *Heligmosomoides* ARI proteins (**Supplementary Fig 1B**) and were named HmixARI.

### Functional properties of ARIs: heparan sulphate binding

For ARIs that contained multiple close homologues in the *Heligmosomoides* genomes, we derived consensus sequences to allow comparison of functional elements (**Figure 2C and Supplementary Figure 2A**). All ARI protein sequences consist of a signal peptide, followed by 3 complement control protein (CCP) domains. Our previous work showed that HbakARI2 binds to HS via a charged patch in the CCP1 domain (17). To assess the presence of the charged patch in CCP1 mediating HS binding, we aligned the CCP1 domains of all ARI proteins to identify charged residues at sites previously implicated in HS interactions (17) (**Figure 3A**). The CCP1 domain from each ARI was then modelled in Alphafold 3 (24) and electrostatic surface charges were calculated. As shown previously (17), HbakARI1 and HbakARI2 CCP1 models show a strong positively charged patch that mediates binding to HS, while HbakARI3 largely lacks this charged site (**Figure 3B-D**). Likewise, modelling of HpolARI1 and HpolARI3 show similar charge patterns to their *H. bakeri* homologues, with HpolARI3 lacking this charged site (**Figure 3E-F**). HpolARI4 shows conservation of charged residues with HbakARI2 (**Figure 3A**) and likewise, electrostatic modelling indicates a similarly charged patch on HpolARI4 CCP1, indicating the potential of HpolARI4 to bind to HS (**Figure 3G**). Finally, HmixARI also shows a positively charged patch on its CCP1 domain (**Figure 3H**), despite multiple polymorphisms within its CCP1 domain compared to HbakARI2 (**Figure 3A**). Therefore, as HpolARI4 has a similar mean electrostatic potential of the surface-exposed residues in its CCP1 domain as HbakARI2 (**Figure 3I**), we proposed that it may also bind to HS in a similar manner to HbakARI2.

**Figure 3:**
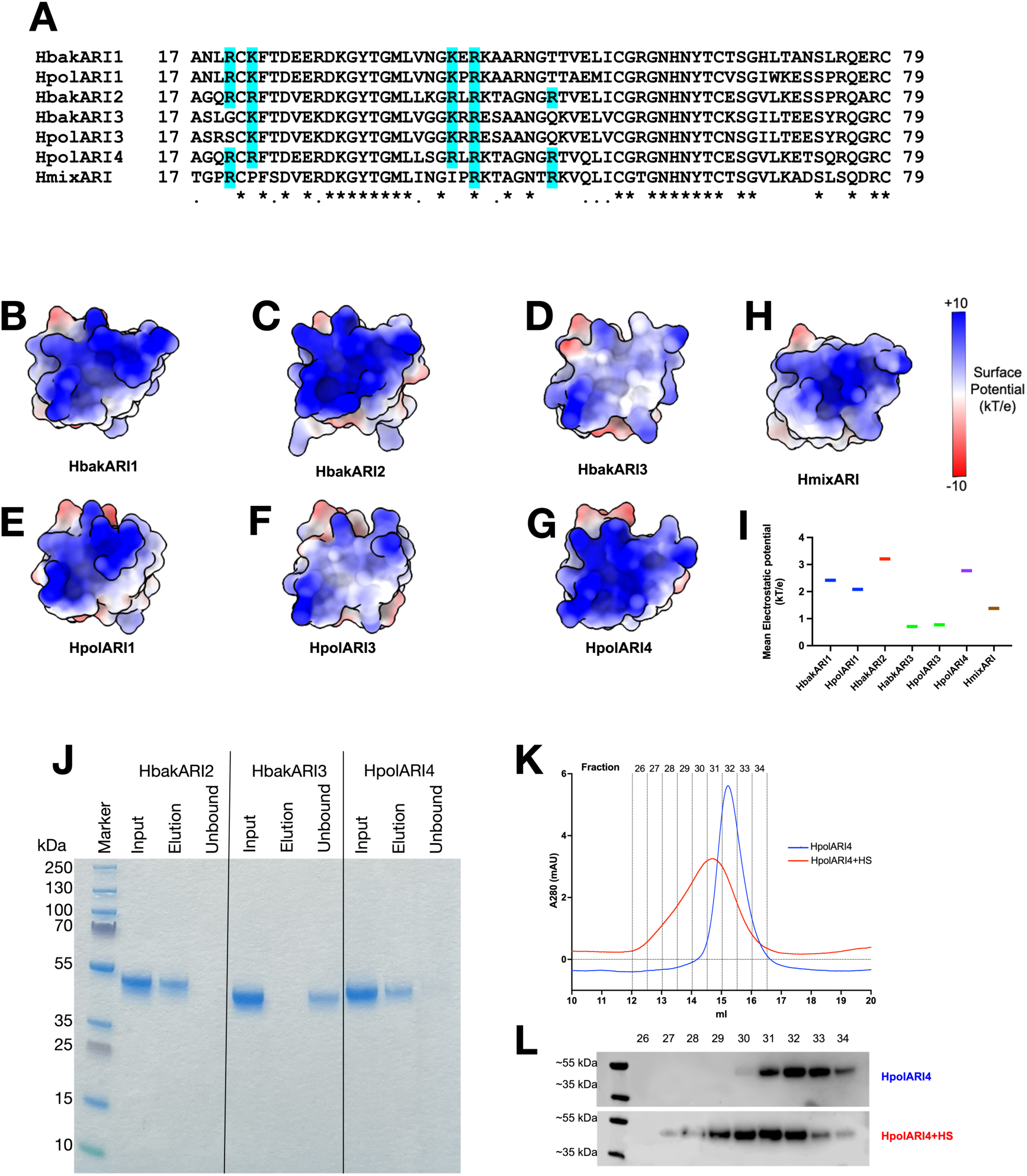
Alignment of CCP1 domains of each ARI, with surface-exposed charged residues implicated in heparan sulphate binding highlighted blue (A). Electrostatic surface rendering of AlphaFold models of the CCP1 domains of ARIs from *H. bakeri* (B, C, D), *H. polygyrus* (E, F, G) and *H. mixtum* (H). Mean electrostatic potential of surface-exposed residues in each CCP1 domain shown in (I). Coomassie gel of HbakARI2, HbakARI3 and HpolARI4 pull down using heparan sulphate coated beads. Input, pull-down and unbound (supernatant) shown (J). Representative of 3 repeats. A280 trace of 50 pg of HpolARI4 +/- 50 pg of heparan sulphate ran on a superdex 200 increase 10/300 GL gel filtration column (K). Representative of 2 repeats. Western blot of gel filtration fractions 26-34 from K stained for His-tag present on HpolARI4 (L).

We expressed HpolARI4 in mammalian cells and purified in the same manner as the previously-expressed HbakARI1, HbakARI2 and HbakARI3 proteins. Recombinant HpolARI4 could bind to heparin-coated beads similarly to HbakARI2 (**Figure 3J**). We confirmed this HS binding by size exclusion chromatography to show a shift of the HpolARI4 elution peak in the presence of HS (**Figure 3K-L**). Together, these data show that conserved features within the CCP1 domain can predict HS binding and identify HpolARI4 as a new HS-binding ARI homologue.

### Functional properties of ARIs: IL-33 binding

The interaction of ARIs with HS has functional consequences for the efficacy and *in vivo* half-life of these proteins, as previously described (17). However, HS is an invariant glycosaminoglycan, and therefore is not subject to host-parasite co-evolutionary pressures, unlike protein targets like IL-33. To assess the affinity and specificity of the ARIs for host IL-33, we focussed on their CCP2 and CCP3 domains (**Figure 4A-B**). Our previous structural studies showed that HbakARI2 binds to *M. musculus* IL-33 via 2 interfaces: interface 1 mediated by residues in the HbakARI2 CCP2 domain, and interface 2 via a large loop in the HbakARI2 CCP3 domain (16). These interfaces are illustrated in **Figure 4C**. We hypothesised that HpolARI4 co-evolved with IL-33 of the parasite’s natural host, *A. sylvaticus*. We therefore modelled the interaction between HpolARI4 and *A. sylvaticus* IL-33, showing a lack of conservation of some residues at interaction site 1 (**Figure 4A, C**). To further investigate a less well-conserved ARI homologue, *H. mixtum* HmixARI was modelled with *M. glareolus* IL-33, and despite multiple polymorphisms in these sequences, a similar structure as that seen for HbakARI2 bound to *M. musculus* IL-33 (16) was predicted (ipTM = 0.78, pTM = 0.8) (**Supplementary Figure 3**). These models support the hypothesis that the novel ARI proteins could adopt conformations compatible with binding to their respective host IL-33 proteins. We next sought to experimentally investigate these potential interactions.

**Figure 4:**
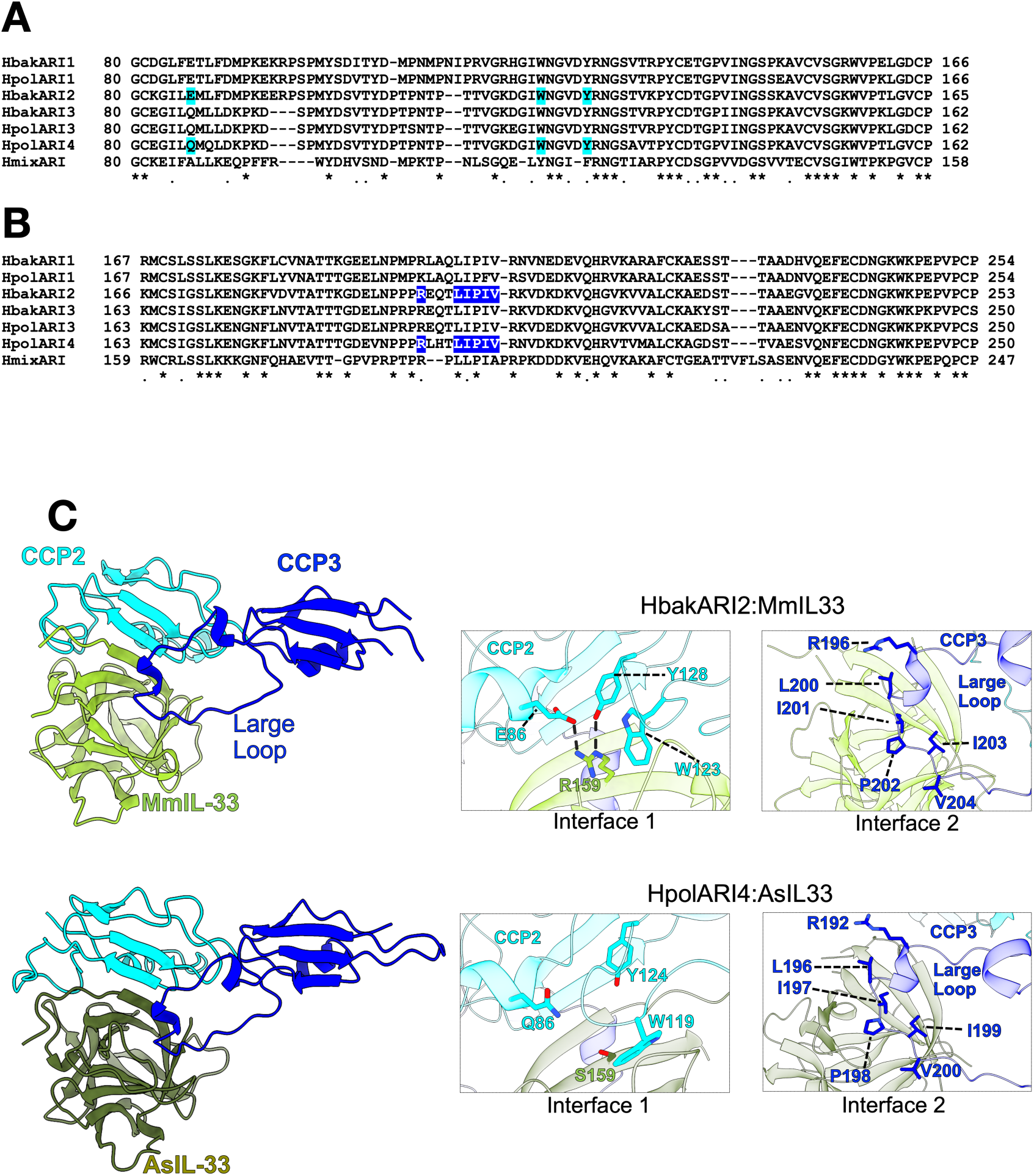
Alignment of consensus *H. bakeri, H. polygyrus* and H. *mixtum* ARI CCP2 (A) and CCP3 (B) domains - residues involved or implicated in IL-33 interfaces are highlighted blue. Zoomed images of *H. bakeri* HbakARI2 - *M. musculus* IL-33 crystal structure (PDB: 8Q5R) interfaces 1 and 2 shown to illustrate interacting residues, with Alphafold 3 model of *H. polygyrus* HpolARI4 bound to A. *sylvaticus* IL-33 shown for comparison (C).

**Supplementary Figure 3:**
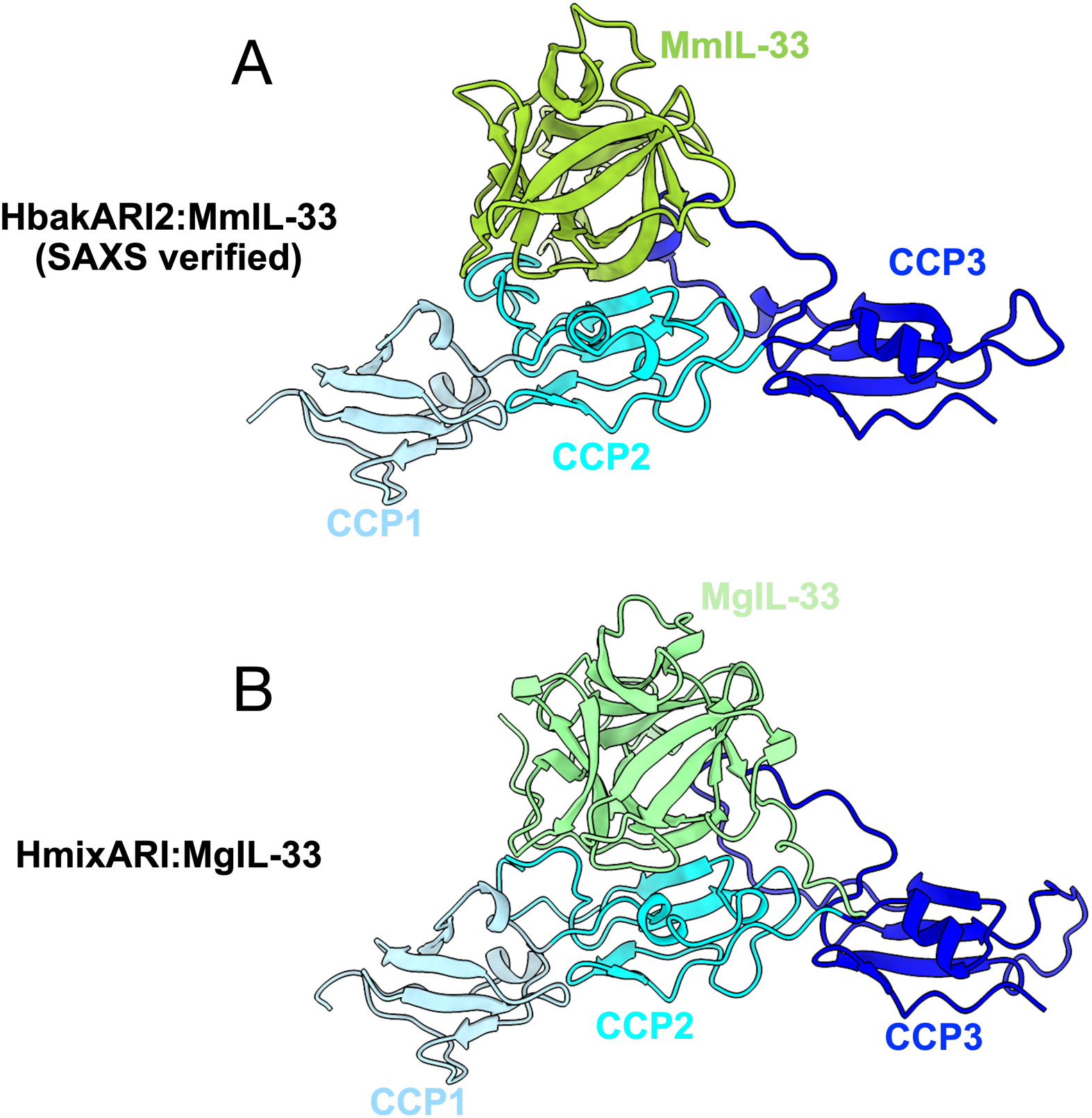
Alphafold 3 models of HbakARI2 with *M. musculus* IL-33 (A) and HmixARI with *M. glareolus* IL-33 (B). ARIs are shown in blue, IL-33s shown in green. HbakARI2: Mm IL-33 model has been verified by small-angle X-ray scattering (SAXS), SASBDB accession code SASDUY6.

The affinities of HbakARI1, HbakARI2 and HbakARI3, and HpolARI4 for *H. sapiens, M. musculus* and *A. sylvaticus* IL-33 proteins were assessed by surface plasmon resonance (**Figure 5**). HbakARI1 showed no binding to human IL-33, while only HbakARI3 showed sub-nanomolar affinity for the human cytokine (**Figure 5A, C, E, G**). Conversely, all ARIs showed sub-nanomolar affinity for *A. sylvaticus* IL-33, with HpolARI4 showing the strongest binding (**Figure 5B, D, F, H**). Finally, while our previous work (14) had established high affinity of binding to *M. musculus* IL-33 by HbakARI1 and HbakARI2 (K_d_ values <0.1 nM), and somewhat lower affinity of HbakARI3 for *M. musculus* IL-33 (K_d_ =0.41 nM), HpolARI4 showed >100-fold lower affinity for *M. musculus* IL-33 (K_d_=78 nM) (**Figure 5I-J**). Therefore, while all HpARIs show moderate to high affinity for *A. sylvaticus* IL-33, only the *H. bakeri*-derived ARIs show strong affinity for *M. musculus* IL-33. By contrast, HbakARI3 showed the broadest binding profile, binding all tested IL-33 proteins with sub-nanomolar affinity.

**Figure 5:**
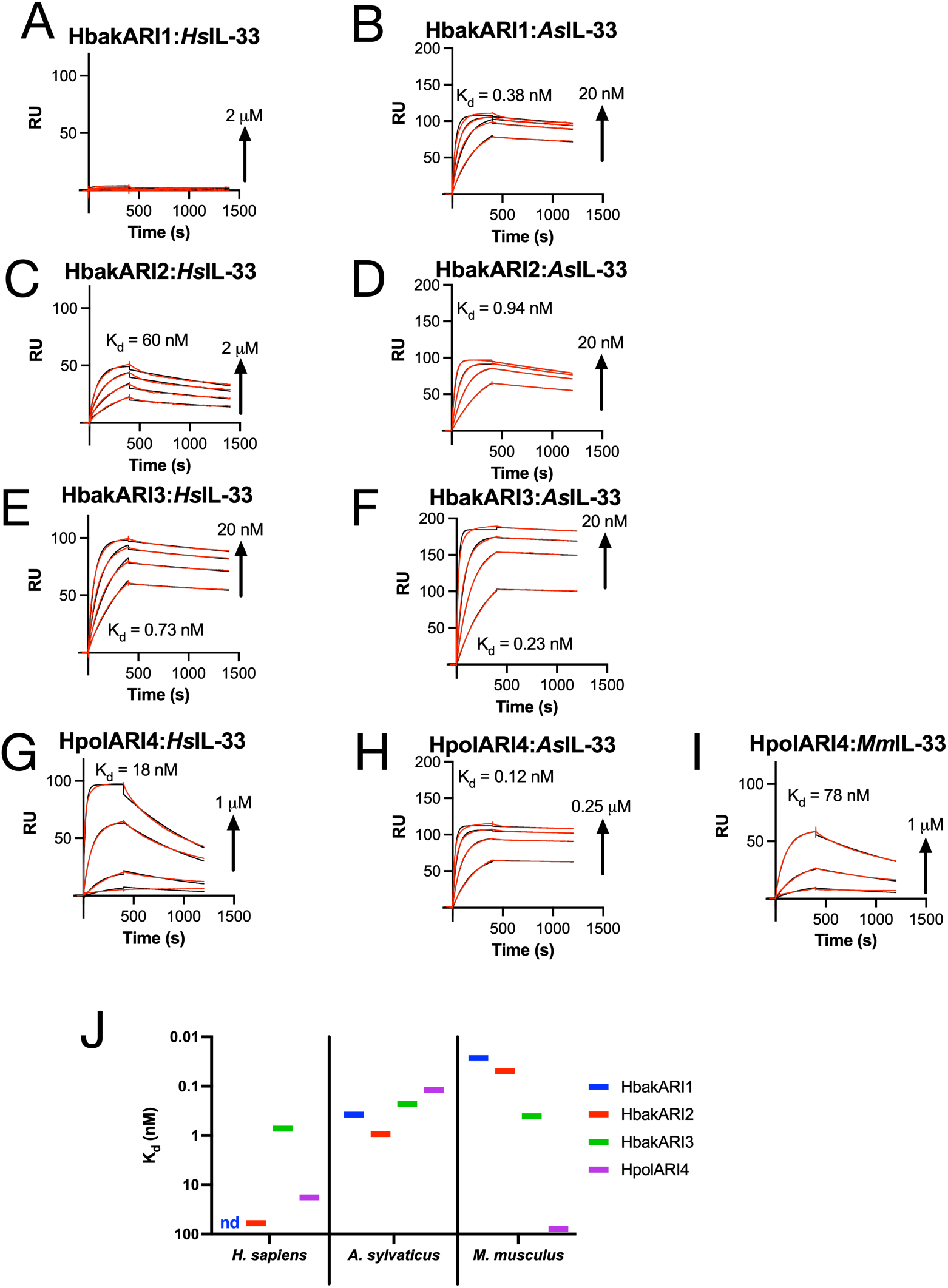
Affinity of ARIs for different species’ IL-33. Surface plasmon resonance was used to measure the affinity for HbakARH (A, B), HbakARI2 (C, D), HbakARI3 (E, F) and HpolARI4 (G, H, I) against *H. sapiens* IL-33 (A, C, E, G), *A. sylvaticus* IL-33 (B, D, F, H) or *M. musculus* IL-33 (I). Affinities for each IL-33 summarised in (J).

We further tested the functional suppressive capacity of each of the ARIs against each IL-33 in *in vitro* assays. Consistent with its high affinity for human IL-33, HbakARI3 suppressed responses to the cytokine-induced responses in human cell assays at sub-nanomolar concentrations (HbakARI3:hIL-33 IC_50_ ∼0.1 nM). Surprisingly, despite only moderate binding affinity for human IL-33, HpolARI4 was still capable of suppressing responses to human IL-33 at high concentrations (HpolARI4:hIL-33 IC_50_ ∼6 nM). By contrast, HbakARI1 and HbakARI2 had no substantial suppressive effect on the response (**Figure 6A**). Collectively, our assays show that HbakARI3 is the most potent suppressor of human IL-33.

**Figure 6:**
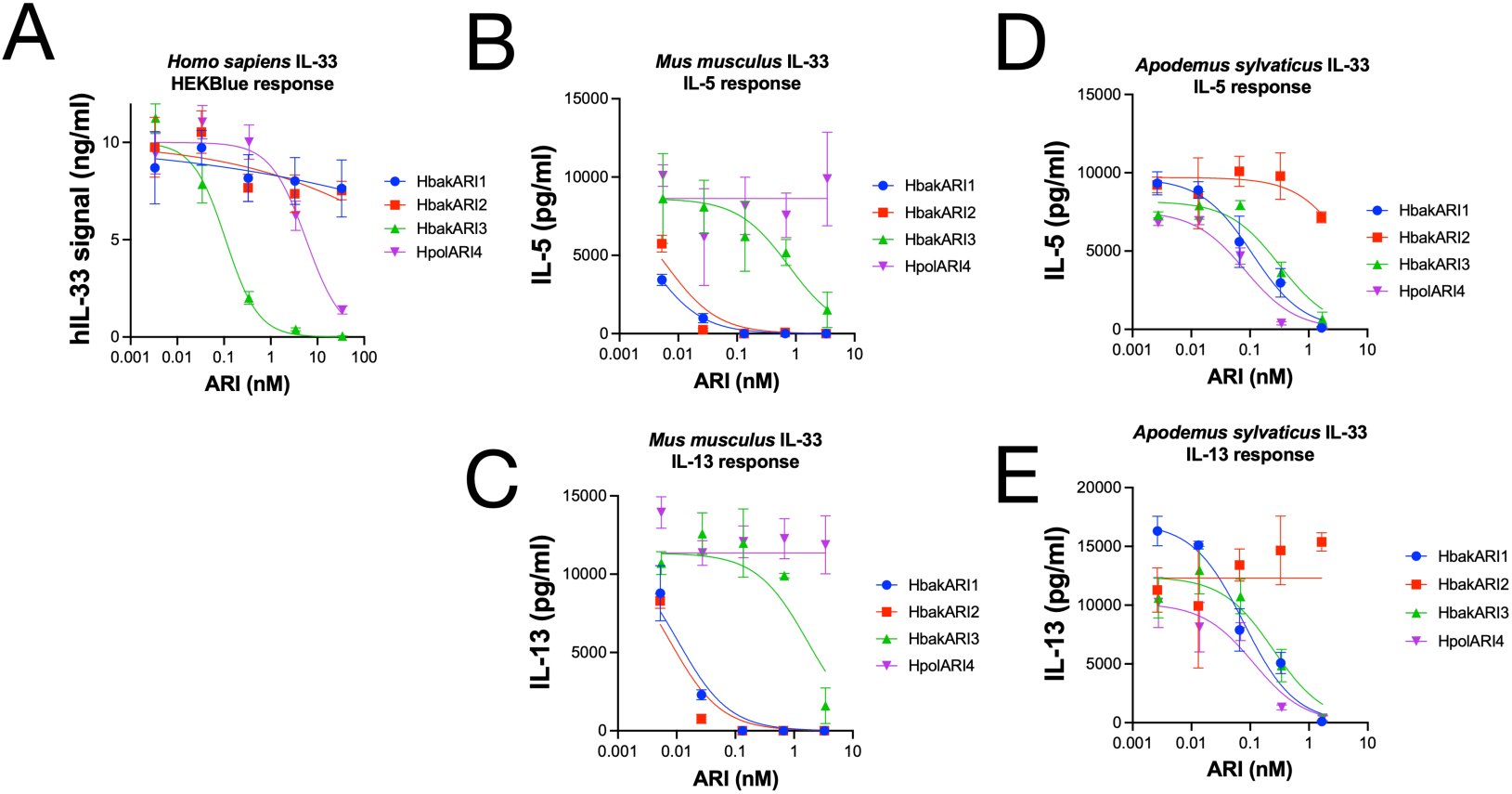
HbakARI2 suppresses lab mouse IL-33 but not wood mouse IL-33. HEKBIue IL-33 reporter cells were stimulated with 10 ng/mL hlL-33 and HbakARIl, HbakARI2, HbakARI3 and HpolARW, and secreted alkaline phosphatase measured 24 h later **(A).** Lab mouse bone marrow was cultured with recombinant *Mus musculus* IL-33 (MmlL-33; 1 ng/mL), IL-2 (10 ng/mL), IL-7, and HbakARIl, HbakARI2, HbakARI3 or HpolARW for 5 days, and **(B)** IL-5 and **(C)** IL-13 in supernatants was assessed by ELISA. Lab mouse bone marrow was cultured with recombinant *Apodemus sylvaticus* IL-33 (AslL-33), IL-2, IL-7, and HbakARIl, HbakARI2, HbakARI3 or HpolARW for 5 days, and **(D)** IL-5 and **(E)** IL-13 in supernatants was assessed by ELISA. Data represent mean +/-SEM data. n=3 biological replicates (i.e. different mouse bone marrow) per group.

To test responses to *M. musculus* and *A. sylvaticus* IL-33s, we turned to rodent cell-based assays. We found that *M. musculus* bone marrow cells were capable of responding to either *M. musculus* or *A. sylvaticus* IL-33, leading to robust IL-5 and IL-13 release. In this assay, HbakARI1 and HbakARI2 were both potently suppressive (at picomolar concentrations) of responses to *M. musculus* IL-33 (both HbakARI1:MmIL-33 and HbakARI2:MmIL-33 IC_50_ values ∼0.01 nM), while HbakARI3 could only suppress responses at the highest concentrations tested, and HpolARI4 showed no effect (**Figure 6B-C**). By contrast, HbakARI1, HbakARI3 and HpolARI4 were all capable of suppressing responses to *A. sylvaticus* IL-33 at sub-nanomolar concentrations (HbakARI1, HbakARI3 and HpolARI4 IC50 values against AsIL-33 ∼0.1-1 nM), while HbakARI2 was ineffective in this assay (**Figure 6D-E**).

Although *A. sylvaticus* IL-33 was effective in inducing responses in *M. musculus* bone marrow cells, there is a mismatch between the source of the IL-33 and the IL-33 receptor in this assay. Therefore, we also stimulated *A. sylvaticus* bone marrow cells with *A. sylvaticus* IL-33. No cross-reactive ELISAs were available for IL-5 or IL-13 to assess the *A. sylvaticus* cellular response, and likewise very few flow cytometry reagents showed cross-reactivity. Nonetheless we found IL-33 stimulation of *A. sylvaticus* bone marrow cells resulted in a significant upregulation in the proportion of ICOS-positive cells in these cultures, and we therefore used this as a readout of IL-33 responsiveness. When ARIs were added to these cultures, we found that HbakARI2 tended to show reduced suppressive capacity of responses, compared to HpolARI4 (**Supplementary Figure 4**), supporting our *M. musculus* results.

**Supplementary Figure 4:**
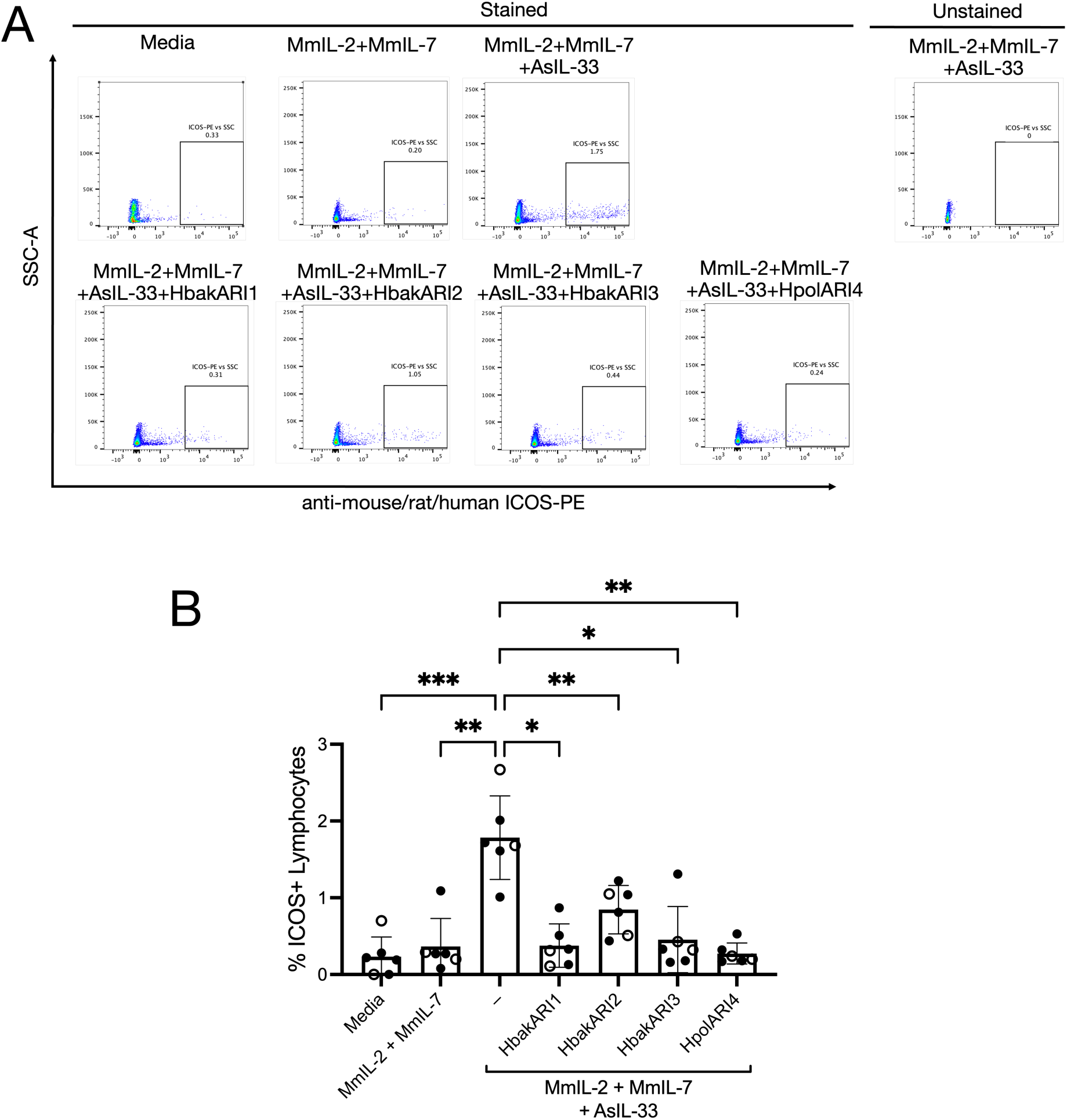
ARI family effects on wood mouse. IL-38-dependent bone marrow responses. Wood mouse bone marrow was cultured with recombinant *Apodemus sylvaticus* IL-33 (AsIL-33), *Mus musculus* IL-2 (MmIL-2), *Mus musculus* IL-7 (MmIL-7), and HbakARI1, HbakARI2, HbakARI3 or HpolARI4 for days, and ICOS staining on live lymphocytes was assessed by flow cytometry. Representative flow staingin shown i **(A)**, refplicates shown in **(B)**. Data shows n=6 biological replicates (i.e. different mouse bone marrw) per group. Males are indicated by solid circle and females indicated by open circle. Standard error of mean shown. Data analysed by one way ANOVA comparing all groups tot positive control,

The crystal structure of the *M. musculus* IL-33-HbakARI2 complex shows two interfaces: interface 1, mediated largely by binding between IL-33 R159 and HbakARI2 CCP2 residues E86, W124 and Y129, and interface 2, mediated by a HbakARI2 CCP3 loop containing R197, L201, I202, P203, I204 and V205, which interacts with a hydrophobic pocket in IL-33 and sterically hinders interactions between IL-33 and its receptor, ST2 (**Figure 4C**) (16). There are notable polymorphisms within these interfaces when comparing the ARIs and IL-33 proteins from different species. Human IL-33 encodes a serine residue at 159 in place of arginine in the *M. musculus* sequence and would therefore lack the charge interactions with HbakARI2 E86. Indeed, when a human IL-33 S159R mutant was expressed, it was demonstrated to have a ∼100-fold higher affinity for HbakARI2 compared to wild-type human IL-33 (16), indicating the importance of this interaction for HpARI2 binding to IL-33. Surprisingly, *A. sylvaticus* IL-33 also contains a serine at the 159 interface site, which is notable as no other rodent species’ IL-33 protein contains an uncharged residue at this site, even those species considerably more distantly related to *M. musculus* than *A. sylvaticus* (**Supplementary Figure 5A-C**). The human IL-33 S159R mutant was also tested for functional suppression by HbakARI1, HbakARI2, and HbakARI3, and while HbakARI1 remained non-suppressive of this mutant, and HbakARI3’s suppressive effects were likewise unchanged, HbakARI2 could only suppress responses to the mutant, but not the wild-type protein (**Supplementary Figure 5D-E**). Therefore, the interface between HbakARI3 and human IL-33, unlike that between HbakARI2 and MmIL-33, may not require a charge interaction. We further hypothesise that HpolARI4 also lacks this charge-based interaction, and allows effective binding to *A. sylvaticus* IL-33.

**Supplementary Figure 5:**
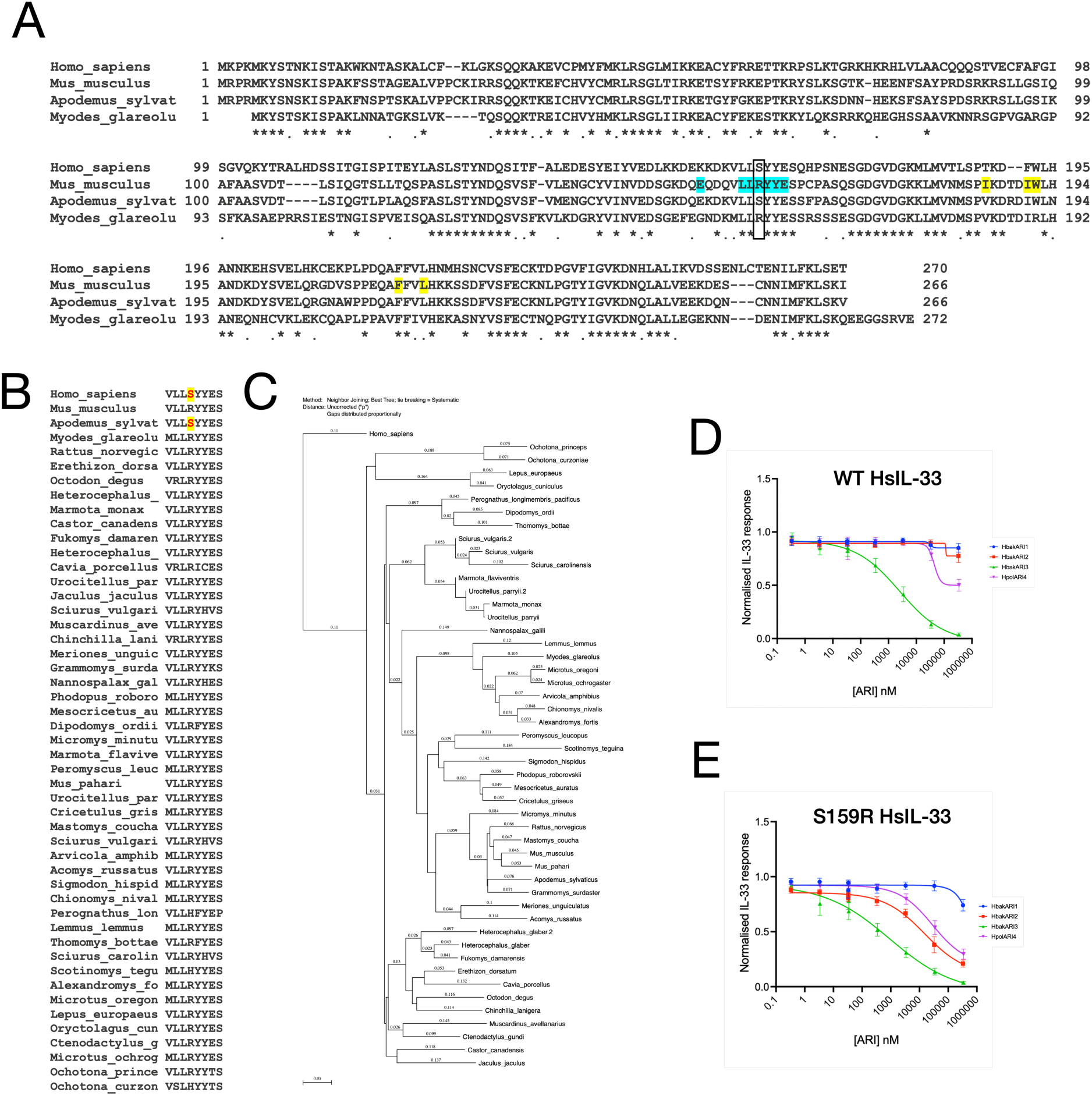
Alignment of IL-33 sequences from human, *M. musculus, A. sylvaticus and M. glareolus* (A). Box indicates *M. musculus* IL-33 R159 binding site for HbakARI2. Interface 1 residues shown in green, interface 2 in yellow. Alignment of IL-33 Interface 1 site in *H. sapiens* and a range of rodent and rabbit species in which IL-33 could be identified, showing that only *H. sapiens* and *A. sylvaticus* encode a serine residue at position 159 (B). Phylogenetic tree of the full-length IL-33 sequences used to produce alignment shown in B, rooted to the *H. sapiens* sequence (C). HEKBIue IL-33 reporter cells were stimulated with (D) *H. sapiens* wild-type IL-33 or (E) *H. sapiens* IL-33 R159S mutant in the presence of HbakARIl, HbakARI2, HbakARI3 or HpolARH. Results are normalised to IL-33-only control and represent 3 experimental repeats.

## Discussion

In this study we identified and characterised a range of heligmosomid ARI family members, including a new member of the family, HpolARI4. We found that the ARIs showed a range of activities, which may reflect their adaptation to their host, with differential affinity for human, *M. musculus* and *A. sylvaticus* IL-33, and differential binding to HS. We found that *H. polygyrus* and *H. bakeri* both encode multiple copies of ARI1, and that these genes are very well conserved within and between species. The HbakARI1 protein binds to HS, is effective at blocking both *M. musculus* and *A. sylvaticus* IL-33, but does not bind to human IL-33. Both *Heligmosomoides* species also encode closely-related ARI3 proteins. HbakARI3 does not bind HS, but is effective against human and *A. sylvaticus* IL-33, but not *M. musculus* IL-33. Finally, while *H. bakeri* encodes HbakARI2, which binds HS and is highly specific for *M. musculus* IL-33, *H. polygyrus* encodes HpolARI4, which binds HS and is specific to *A. sylvaticus* IL-33. Therefore, these parasites have evolved immunomodulators that are highly specific to their natural host targets.

The specificity of these immunomodulators has implications for parasite-host co-evolution, and parasite vaccine development. The radical alteration in efficacy of ARIs against *M. musculus* versus *A. sylvaticus* IL-33 implies a Red Queen hypothesis interaction (2), where both parasite and host are both under strong selection pressure. In particular, it is noteworthy that at a critical binding site (R159) in the IL-33 sequence, *A. sylvaticus* has an R>S polymorphism, which is shared with *H. sapiens* but is not found in any other rodent IL-33 sequences analysed. This polymorphism could be the result of selection pressure from the effects of parasite-derived ARI proteins. A parallel example of a single, functionally critical residue shaping host infection is provided by the *Plasmodium falciparum* reticulocyte-binding protein homologue 5 (RH5), which is required for erythrocyte invasion via basigin. Ancestral RH5 could bind both gorilla and human basigin but a single H200Y substitution abolishes gorilla binding while preserving human basigin binding (25). In this context, it may be useful to consider the ARI homologue in *H. mixtum*, which we propose to bind to IL-33 from its host *M. glareolus*, and shows a notable lack of conservation at the interface 1 binding site (49-55% identity with other ARIs across the whole protein sequence, with only 26-31% identity in the interface 1 region), again implying that these immunomodulators are under strong selection pressure.

The lack of defined host-pathogen interaction proteins has resulted in less interest in structure-function relationships within nematode parasite co-evolution than in other pathogens such as viruses, bacteria or unicellular organisms. For instance, the evolution of the MERS-CoV spike protein defines host specificity as it mediates entry via interaction with host ACE-2 receptor, and a 5 amino acid polymorphism in hamster ACE-2 renders this species resistant to infection (26). Similarly, the immunomodulatory bacterial molecule, *Neisseria gonorrhoeae* factor H binding protein (fHBP), binds to host complement factor H (CFH), and a single-residue polymorphism (N1203R) in chimpanzee CFH renders them resistant to both fHBP binding and infection by the bacteria (27). These studies clearly underline the importance of specificity in host-pathogen molecular interactions, especially in the context that both coronavirus spike proteins and *Neisseria* fHBP are effective as vaccine antigens, due to raising of antibody responses which block interaction between the pathogen and host proteins (2). However, these interactions are complex and efficacy cannot always be correlated with conservation of binding sites: in a study of 3 fHBP variants, critical binding site residues were found to be mutated in each case, but compensated for by forming new interactions, resulting in each variant having a similar affinity and efficacy for the host target (28). Thus, pathogen-host interactions must always be empirically confirmed. This is especially important as these immunomodulatory host interaction proteins are developed as vaccine candidates: our recent work shows that only HbakARI2 (and not HbakARI1 or HbakARI3) vaccination was effective in protection against *H. bakeri* infection (18), and the results shown here lead us to propose that HpolARI4 would be the most effective vaccine antigen in *H. polygyrus* infection of *A. sylvaticus*. Likewise, in other parasite-host combinations (e.g. the *Trichuris* family of p43-IL-13 binding proteins (29)), the correct immunomodulatory vaccine antigen must be chosen for effective protection against infection.

The variety of the ARI family also leads to questions about the evolution and role of these proteins. It is notable that both *H. polygyrus* and *H. bakeri* genomes contain multiple copies of ARI1 on chromosome V, which appear to have been produced by gene duplication events. Gene duplication is often associated with very high expression (30) or duplication followed by neofunctionalization, potentially leading to immune escape (31). Neither explanation appears to fit the HbakARI1 situation as it is expressed at a far lower level than HbakARI2 during the first week of infection (14) (where immunomodulation appears to be most active (18)), the multiple ARI1 copies are well-conserved (therefore do not appear to have undergone neofunctionalization), and the HbakARI1 protein is ineffective as a vaccine antigen (18) (therefore it does not seem dominant in the host-parasite interaction). Furthermore, the duplication events in *H. bakeri* appear to be more recent than in *H. polygyrus*, as the HbakARI1 copies are very well-conserved in sequence and orientation, while the HpolARI1 copies show greater deviation from one another. In *in vivo M. musculus* experiments, HbakARI1 showed reduced suppressive capacity compared to HbakARI2 (14) (in contrast to their very similar effects *in vitro* shown here), while previous assessment of the HS binding activity of HbakARI1 showed it had reduced HS binding capacity compared to HbakARI2 (17), and therefore may diffuse more freely from the site of deposition. Antibody responses to HbakARI1 during infection could therefore preferentially bind this poorly effective protein at sites distal to the parasite, allowing HbakARI2 to impede IL-33 responses at sites proximal to the parasite without being neutralised by antibodies.

Similar to ARI1, ARI3 is well-conserved between *H. polygyrus* and *H. bakeri*. It is the most promiscuous of the ARIs, with HbakARI3 showing sub-nanomolar affinity for all three IL-33 species tested, and showing suppressive activity against each in vitro. However, our previous work showed that HbakARI3 can stabilise *M. musculus* IL-33 *in vivo*, and after release of endogenous IL-33 from necrotic epithelial cells in culture (14). It is unclear why HbakARI3 can have opposing effects on recombinant versus endogenously-released IL-33, however this further highlights the need for *in vivo* testing of these immunomodulatory proteins.

We were surprised that a close homologue of HbakARI2 could not be identified in the *H. polygyrus* genome, but instead identified a new member of the family (HpolARI4) that encoded elements of both HbakARI2 (CCP1 domain homology and HS binding) and HbakARI3 (CCP2+3 domain homology and suppression of *A. sylvaticus* and human IL-33). Due to its localisation on chromosome III, at a similar site to HbakARI2 in *H. bakeri,* it is likely that HbakARI2 and HpolARI4 evolved from the same ancestral protein, as *Heligmosomoides* species adapted to *M. musculus* and *A. sylvaticus* hosts. We furthermore hypothesise that HpolARI4 would be the most effective vaccine candidate in *H. polygyrus - A. sylvaticus* infections, similarly to the efficacy of the HbakARI2 vaccine in *H. bakeri* – *M. musculus* infections (18).

Finally, this study assessed the suppressive capacity of the ARIs against human IL-33 and found that HbakARI3 had the strongest suppressive effect against human IL-33 *in vivo* and *in vitro*. Despite HpolARI4 and HbakARI3 sharing many features of their CCP2/3 IL-33 binding domains, HbakARI3 showed a clear advantage over HpolARI4 in suppression of responses to human IL-33 in vitro. However, as HbakARI3 lacks HS binding, it has a short half-life in vivo (17). To address this, the HbakARI2:3 fusion protein used here in vivo may be the most successful in suppression of human IL-33 responses, and could form the basis for design of more effective protein inhibitors of IL-33 for use in human disease.

## Conclusions

This study shows that nematode immunomodulatory proteins adapt to show fine scale specificity for their host species. Detailed understanding of these parasite-host interactions will not only enable a deeper understanding of their co-evolutionary relationships but also allow development of effective vaccination and treatment regimes.

## Methods

### Animals

C57BL/6JCrl mice were purchased from Charles River, UK. Human IL-33 transgenic mice (hIL-33^+/+^mIL-33^−/−^, “hIL-33tg”) on a C57BL/6J background have been described previously (21). Mice were used at 8-12 weeks old, with both sexes used unless otherwise indicated.

Wood mice (A. sylvaticus) are maintained in standard laboratory conditions at the University of Edinburgh. These animals have been maintained in captivity for many generations but retain both genetic and microbiome diversity. Mice were co-housed in single sex groups in ventilated cages with ad libitum food and water. Individuals of both sexes were used, and within sexes and replicates, siblings were used. Wood mice were between 10-11 months at time of tissue collection.

### Protein expression

ARI protein expression was carried out as previously described (14). Briefly, plasmid constructs expressing the ARIs, or mutants thereof, (with C-terminal c-myc and 6His tags) were transfected into Expi293 cells using the Expifectamine transfection kit, with addition of enhancers 24 h later, following manufacturer’s instructions (ThermoFisher). Culture supernatants were collected 4-7 days later, and proteins purified using nickel column purification. Proteins were dialysed into PBS and filter sterilised.

*M. musculus* and Human IL-33s were expressed and purified as described in previous publications (16). A similar expression strategy was used to obtain highly pure *A. sylvaticus* IL-33. Briefly, the *A. sylvaticus* IL-33 gene (NCBI XP_052043329, residues 109-270) was cloned into pET28a and expressed in BL21-DE3 as a TEV protease-cleavable N-terminal 6His-tag protein. Expressed protein was purified using Ni-NTA chromatography, and eluted fractions were pooled and concentrated to 3 mg/ml. Concentrated protein was then injected into a S75 10/300 gel filtration column equilibrated with 20 mM HEPES pH 7.2 and 150 mM NaCl to obtain pure and homogenous protein for functional studies.

All proteins were assessed for the absence of endotoxin using the HEK-Blue LPS Detection kit 2 (InvivoGen). All proteins contained <0.1 EU LPS per μg of protein.

### Alternaria model

Female hIL-33tg mice were briefly anaesthetised using inhalational isoflurane. *Alternaria alternata* allergen (50 μg per mouse) was administered intranasally in 50 μl of PBS, alone or in the presence of 10 μg of HbakARI1, HbakARI2, HbakARI3 or HbakARI2:3. Mice were culled either 30 min or 24 h later by overdose of injectional anaesthetic, followed by severing of a major vessel. BAL cells and fluid were collected via 3 washes with 0.5 ml of ice-cold PBS using an 18G needle. Lungs were dissected, and the right lobes were taken for single cell preparations of cells by digesting minced tissue in 1ml PBS (+Mg^2+^ and Ca^2+^) containing Liberase TL (2 U/ml; Roche) and DNAse1 (160 U/ml; Sigma) for 35 min, shaking at 200 rpm at 37°C. The digest was then stopped with 5 mL ice-cold complete RPMI (Gibco) and crushed through a 70 μm nylon filter (Greiner). Red blood cells were lysed in ACK lysis buffer (Gibco), then resuspended in PBS for counting.

### Flow cytometry

BAL and lung cells were stained for flow cytometry by first resuspending in 1 x PBS containing Fixable Blue Dead Cell Stain Kit (1:1,000, Invitrogen for 15 min in the dark at 4°C. Cells were centrifuged (400 g, 5 min, 4°C) and resuspended in FACS buffer (PBS with 0.5% BSA and 0.05% sodium azide) containing purified anti-mouse CD16/32 (Clone: 93, BioLegend, 1:50) and incubated at 4°C for 20 min. Cells were then washed with FACS buffer, and stained for flow cytometry at 4°C for 20 min using combinations of the following antibodies: CD45-AF700 (clone: 30-F11, 1:200); CD3-FITC (clone: 145-2C11, 1:200); CD5-FITC (clone: 53–7.3, 1:200); CD11b-FITC or -Pacific Blue (clone: M1/70, 1:200); CD11c-AF647 (clone: N418, 1:200); CD25-BrilliantViolet 650 (clone: PC61, 1:200); CD4-PE-Dazzle (clone: RM4-5, 1:200); Gr1-FITC (clone: RB6-8C5, 1:200); CD19-FITC (clone: 6D5,1:200); ICOS-PE (clone: C398.4A, 1:200); CD5-FITC (clone: 53–7.3, 1:200); ST2-APC (clone: RMST2-2, 1:100); or CD49b-FITC (clone: DX5, 1:200) from Invitrogen, or Siglec-F-PE (clone: REA798,1:200) from Miltenyi Biotec. For transcription factor staining, cells were then incubated overnight at 4°C in FoxP3 fixation/permeabilisation buffer (ThermoFisher), then resuspended in 1 x permeabilisation buffer containing GATA3-PE (clone: TWAJ, 1:50) for 30 min at 4°C. Samples were acquired on a LSR Fortessa (BD), and analysed with FlowJo v10.9 (Waters Biosciences). Cells were gated on live singlets, and then eosinophils were gated on SiglecF^hi^CD11c^−^CD45^+^ cells and ILC2s were gated on GATA3^+^CD3^−^CD5^−^CD19^−^GR1^−^CD11b^−^ CD4^−^CD45^+^ cells.

### Surface Plasmon Resonance

Purified ARI family members in 1XPBS were biotinylated by mixing 50 mM of protein with 100 mM of EZ-NHS-Biotin (Thermofisher Scientific) followed by incubation for 2 h on ice. Excess biotin was removed using a PD-5 column 20 mM Tris-Cl pH, 8.0 and 150 mM NaCl. Experiments were performed at 25°C on a Biacore T 200 instrument using Biotin Capture kit (Cytiva) in a buffer containing 20 mM Tris-Cl, pH 8.0, 150 mM NaCl, 0.05 % Tween-20 and 1 mg/ml salmon sperm DNA. 300-400 RU of each biotinylated ARI were immobilised on different flow cells. The binding measurements were performed at a flow rate of 40 μl/min by injecting two-fold concentration series of various IL-33. The data was processed using BIA evaluation software version 1.0 (BIAcore, GE Healthcare).

Response curves were double referenced by subtracting the signal from the reference cell and averaged blank injection. Each experiment was performed twice.

### Heparin-agarose pull-down assays

Heparin-agarose beads (Sigma) were incubated with 10 μg of HbakARI2, HbakARI3 or HpolARI4 proteins in 100 μl of PBS+0.02% Tween 20 and rotated for 30 min at room temperature. Beads were then spun down and the supernatant collected, and washed three times with PBS+0.02% Tween 20. Bound proteins were eluted in 1X Loading Sample Buffer (ThermoFisher) containing 5% β-mercaptoethanol, heated to 70°C for 5 min and ran on a 4–12% NuPAGE precast gel (ThermoFisher), prior to visualisation using InstantBlue Coomassie Stain (Abcam) following the manufacturer’s instructions.

### Gel filtration

HpolARI4 protein (50 μg) was incubated alone, or in the presence of 50 μg of HS (HS sodium salt from bovine kidney, Sigma) in PBS for 30 min at room temperature, then applied to a Superdex 200 Increase 10/300 GL gel filtration column (Cytiva) and elution profile detected by monitoring of UV A280 nm absorbance. Collected fractions were ran under reducing conditions on a 4-12% NuPAGE precast gel (ThermoFisher), transferred to nitrocellulose membrane and blocked using 1% BSA in TBS. Membrane was probed with 6x-His Tag rat monoclonal antibody (Invitrogen) overnight at 4°C. The membrane was washed with 1X TBST 5 times then incubated with goat polyclonal anti-rat IgG-HRP (Abcam) for 1hr, then washes repeated. Bands were visualised using chemifluorescence substrate (ThermoFisher Scientific) and captured on a LiCor Odysses Fc.

### M. musculus and A. sylvaticus bone marrow assays

Bone marrow was collected from *M. musculus* C57BL/6J mice or *A. sylvaticus* wood mice by flushing the tibias and femurs with RPMI 1640 using a 23g needle and cell suspensions were passed through a 70 μm strainer. Cells were resuspended in ACK lysis buffer (Gibco) for 5 min at room temperature, then washed and resuspended in RPMI 1640 medium supplemented with 10% fetal bovine serum, 10% fetal bovine serum (ThermoFisher), 2 mM GlutaMAX, 100 IU/mL penicillin and 100 μg/mL streptomycin (Gibco). Cells were cultured in 200 μl volumes in round bottom, 96-well plates at a density of 2.5 x 10^6^ cells/ml.

Recombinant *M. musculus* IL-2 and IL-7 (Biolegend) were added to final concentrations of 10 ng/ml each. Where indicated, samples also included 1 ng/mL of *M. musculus* or *A. sylvaticus* IL-33 and ARI protein at concentrations indicated.

### ELISA

BAL supernatants and bone marrow culture supernatants were assessed for IL-5 and IL-13 concentrations using uncoated ELISA kits (ThermoFisher), following manufacturer’s instructions. Human IL-33 concentration was also assessed in BAL supernatants by IL-33 Duoset ELISA (RCD Systems).

### HEKBlue human IL-33 assay

HEK-Blue IL-33 reporter cells (InvivoGen: hkb-hil33) were used to detect IL-33 signalling via NF-κΒ and AP-1 dependent secretion of alkaline phosphatase. Cells were maintained at 37°C and 5% CO_2_ in Dulbecco’s modified Eagle’s medium supplemented with 10% fetal bovine serum (ThermoFisher), 2 mM GlutaMAX, 100 μg/mL Normocin (InvivoGen) and 1x HEK-Blue selection (InvivoGen) according to the manufacturer’s instructions.

Cells were seeded into 96-well plates at a density of 5 x 10^4^ cells per well and stimulated for 20 hours with 10 ng/mL human IL-33 and, where indicated, ARI protein serially diluted 10-fold. 20 μL of supernatant from each well was combined with 180 μL of Quanti-Blue solution. After 90 minutes of incubation at 37°C, absorbance at 620 nm was measured.

### Statistics

Statistical data were analysed using GraphPad Prism v11.0.2. Independent groups were compared by one-way analysis of variance (ANOVA) with Dunnett’s post-test was used. Error bars show standard error of mean. ****=p<0.0001, ***=p<0.001, **=p<0.01, *=p<0.05, ns = not significant (p>0.05).

## Declarations

### Ethics approval

Mouse accommodation and procedures were performed under UK Home Office licenses with institutional oversight performed by qualified veterinarians. Experiments were approved by the local Welfare and Ethical use of Animals in Research committee (WEC), and conform to relevant regulatory standards.

### Consent for publication

Not applicable

### Availability of data and materials

The datasets used and/or analysed during the current study are available from the corresponding author on reasonable request.

### Author’s contributions

Conception or design of the study: ABP, LW, ESC, MKH, HJM. Acquisition of data: BW, AJ, FC, AO, OCAF, SD, FAJD, LG, DJS, LW, HJM. Analysis of data: BW, AJ, LS, HJM. Interpretation of data: BW, AJ, LS, MKH, HJM. Drafting of manuscript: BW, AJ, ABP, LW, MKH, HJM.

## Acknowledgements

Not applicable

